# Development of the P300 from childhood to adulthood: a multimodal EEG and MRI study

**DOI:** 10.1101/304832

**Authors:** Knut Overbye, Rene J. Huster, Kristine B. Walhovd, Anders M. Fjell, Christian K. Tamnes

**Author notes:** Correspondence to: Knut Overbye, Department of Psychology, University of Oslo, PO Box 1094 Blindern, 0317 Oslo, Norway.

## Abstract

Maturation of attentional processes is central to cognitive development. The electrophysiological P300 is associated with rapid allocation of attention, and bridges stimulus and response processing. P300 is among the most studied and robust electrophysiological components, but how different subcomponents of the P300 develop from childhood to adulthood and relate to structural properties of the cerebral cortex is not well understood. We investigated age-related differences in both early visual and P300 components, and how individual differences in these components related to cortical structure in a cross-sectional sample of participants 8-19 years (n=86). Participants completed a three-stimulus visual oddball task while high-density EEG was recorded. Cortical surface area and thickness were estimated from T1-weighthed MRI. Group-level blind source separation of the EEG data identified two P300-like components, a fronto-central P300 and a parietal P300, as well as a component reflecting N1 and P2. Differences in activity across age were found for the parietal P300, N1 and P2, with the parietal P300 showing stronger activity for older participants, while N1 and P2 were stronger for younger participants. Stronger P300 components were positively associated with task performance, independently of age, while negative associations were found for P2 strength. Parietal P300 strength was age-independently associated with larger surface area in a region in left lateral inferior temporal cortex. We suggest that the age differences in component strength reflect development of attentional mechanisms, with increased brain responses to task-relevant stimuli representing an increasing ability to focus on relevant information and to respond accurately and efficiently.

## INTRODUCTION

From childhood to adulthood, the brain undergoes both complex changes in functional activation patterns and multifaceted regional changes in structural architecture (Blakemore, 2012; Jernigan, Baaré, Stiles, & Madsen, 2011; Segalowitz, Santesso, & Jetha, 2010). These maturational processes are associated with improved cognitive performance, where attentional development plays a key role (Rueda, 2013). The ability to rapidly allocate attention can in part be indexed by the electrophysiological component P300 (Polich, 2007). Previous developmental studies of P300 components have yielded inconsistent findings on the effects of age. Also, there are no studies on the relationships between P300 and brain structure during development. In this study we examined age-related differences in the strength of the P300 and the early visual components that precede it, and how these components are related to both task performance and structural properties of the cerebral cortex.

The P300 is an endogenous positive-going event-related potential elicited in tasks requiring stimulus discrimination, regardless of sensory modality (Polich, 2007; Sutton, Braren, Zubin, & John, 1965; Twomey, Murphy, Kelly, & O’connell, 2015). It is produced when attending to a stimulus and is often interpreted as the first major component of controlled attention (S. Segalowitz & Davies, 2004). However, some argue that earlier components might also reflect aspects of top-down attention, such as the visual N1 (Enge, Fleischhauer, Lesch & Strobel, 2011) and P2 (Sugimoto & Katayama, 2013). Following these early visual components, the P300 peaks sometime after 300 milliseconds (ms) (Linden, 2005; Polich, 2007). The P300 is thought to reflect several overlapping cognitive processes and has been suggested to represent updating of stimulus representations (Donchin, 1981; Polich, 2007), activation of a global conscious workspace (Dehaene, Sergent, & Changeux, 2003), and inhibition of extraneous brain activation to facilitate memory processing (Polich, 2004). It is not solely related to the stimulus or the response to the stimulus but is assumed to be an integrative component bridging the two (Verleger, 2010). Moreover, there are also several lines of research indicating that the P300 can be broken down into various subcomponents such as the P3a and P3b. In addition, principal component analysis (PCA) or independent component analysis (ICA) can be used to decompose the P300 (Eichele, Rachakonda, Brakedal, Eikeland, & Calhoun, 2011; van Dinteren, Huster, Jongsma, Kessels, & Arns, 2017).

Changes in absolute P300 amplitude from childhood to adulthood might reflect improved allocation of cognitive resources (van Dinteren, Arns, Jongsma, & Kessels, 2014b). The P300 signal’s association with executive functions such as attentional control and inhibition, makes it a suitable target for studying functional brain development over adolescence, a period when these cognitive functions are still developing rapidly (Crone & Steinbeis, 2017; Huizinga, Dolan, & van der Molen, 2006). Several cross-sectional developmental studies have examined age-related differences in P300 components, and the oddball paradigm is the most commonly used experimental task (Davies, Segalowitz, & Gavin, 2004; DeBoer, Scott, & Nelson, 2005). These studies generally suggest that P300 amplitudes increase with age, as shown in a meta-analysis and two large cross-sectional studies on auditory P300 components by van Dinteren and colleagues (van Dinteren, Arns, Jongsma, & Kessels, 2014a; van Dinteren et al., 2014b) (see also studies on auditory or visual components by Brinkman & Stauder, 2008; Johnstone et al., 2007; Mullis, Holcomb, Diner, & Dykman, 1985). However, for the P300 measured in visual paradigms, the opposite has also sometimes been reported; namely a decrease in amplitude with increasing age (Berman, Friedman, & Cramer, 1990; Davies et al., 2004; Mahajan & McArthur, 2015; Oades, Dmittmann-Balcar, & Zerbin, 1997; Stige, Fjell, Smith, Lindgren, & Walhovd, 2007). It has thus been suggested that the P300’s development is modality specific (Segalowitz et al., 2010). All studies mentioned used some variant of extracting EEG amplitudes from set time windows. However, since the P300 potential is driven by signals of multiple temporally and spatially overlapping sources (R van Dinteren et al., 2017), P300 potential amplitudes may sometimes be difficult to interpret (Huster, Plis, & Calhoun, 2015). This might be further complicated by possible developmental shifts in P300 source location and latency (Polich, 2007; Segalowitz et al., 2010). In the present study, we aimed to address these complicating phenomena by decomposing the signals using a blind source separation algorithm that is robust to topography and latency differences across individuals (Huster et al., 2015), and thus provide a clearer picture of age-related differences in P300 components, as well as early visual components, from childhood to adulthood. In contrast to the P300, early visual components such as the N1 and P2 have consistently been shown to decrease in amplitude through adolescence (Itier & Taylor, 2004; Tomé, Barbosa, Nowak, & Marques-Teixeira; 2015, Ponton, Eggermont, Kwong, & Don, 2000).

Multiple brain regions are known to contribute to P300 generation (Friedman, 2003). Source localization studies indicate that multiple frontal and parietal brain regions (Bocquillon et al., 2011; Wronka, Kaiser, & Coenen, 2012), as well as temporal and parieto-occipital cortices are involved (Halgren et al., 1995; Mahajan & McArthur, 2015; Smith et al., 1990). P3a sources seem to be situated more anteriorly than those of the P3b (Bocquillon et al., 2011; Wronka et al., 2012). A previous study of adults found higher P300 amplitudes to be related to greater cortical thickness in temporo-parietal and orbitofrontal cortices, but only in older adults (Fjell, Walhovd, Fischl, & Reinvang, 2007). No previous studies have investigated the relationships between the P300 components and brain structure in development.

In the present study, we investigated age-related differences in both early visual and P300 components, and how individual differences in these components relate to task performance and cortical surface area and thickness. We analyzed electroencephalography (EEG) recorded during a three-stimulus visual oddball paradigm and structural magnetic resonance imaging (MRI) data from 86 children and adolescents (ranging from 8 to 19 years). Group-level blind source separation was applied to the EEG data to estimate independent components of brain activity, and FreeSurfer was used to estimate vertex-wise cortical surface area and thickness. We expected the EEG decomposition to result in one or more components representing the P300. In line with recent large studies and a meta-analysis that have shown P300 amplitudes to increase until young adulthood (van Dinteren et al., 2014a, 2014b), we expect the P300 components to be stronger for older participants, reflecting age-related increases in brain responses to task-relevant stimuli. However, negative age-relationships for our visual P300 components would lend credence to the hypothesis of modality specific developmental patterns, as this has been reported in some studies using visual paradigms (Segalowitz et al., 2010). We also expect task performance to improve with age, and to be positively correlated with the strength of the P300 components. In the age-range studied, cortical thickness decreases substantially, while surface area increases during childhood and is relatively stable during adolescence (Amlien et al., 2014; Brown et al., 2012; Raznahan et al., 2011; Tamnes et al., 2017; Vijayakumar et al., 2016). Based on this, we tentatively hypothesize that stronger P300 components would be related to thinner cortex and larger cortical surface area in regions associated with P300 generation. Due to the wide cortical distribution of possible sources of the P300 components, and the sparse literature on relationships between P300 and cortical structure, these analyses were explorative and performed on a vertex-wise level across the cortical surface. Relationships between P300 strength and both task performance and cortical structure are predicted to hold when controlling for age, reflective of individual differences in the phase of functional brain development.

## MATERIALS AND METHODS

### Participants

Children and adolescents between 8 and 19 years of age were recruited to the research project *Neurocognitive Development* (Tamnes et al., 2010; Østby et al., 2009) through newspaper advertisements, and local schools and workplaces. Written informed consent was provided by all participants over 12 years, as well as from a parent or guardian of participants younger than 18 years. Oral informed assent was obtained from participants younger than 12 years. The study was approved by a Norwegian regional committee for medical and health research ethics. Participants aged 16 years or older and a parent completed standardized health interviews regarding each participant. Exclusion criteria included premature birth, a history of injury or disease known to affect central nervous system (CNS) function, ongoing treatment for a mental disorder, use of psychoactive drugs known to affect CNS functioning, and MRI contraindications. Participants were also required to be right-handed, fluent Norwegian speakers, and to have normal or corrected-to-normal hearing and vision. A total of 100 children and adolescents fulfilled these criteria, completed the EEG recording and MRI scanning described below, and were by a neuroradiologist deemed free of significant brain injuries or conditions. Data sets from 12 participants were excluded from analyses due to noisy EEG recordings. Two further subjects were excluded due to behavioral test criteria (described in the next section). The final sample for our analyses thus included 86 participants (43 female) aged 8.2-19.7 years (mean = 14.4, SD = 3.5). Mean ages for males (14.1 years, SD = 3.6) and females (14.7 years, SD = 3.4) were not significantly different (t = 0.76, p =. 46). Mean IQ, as estimated by the four-subtest form of the Wechsler Abbreviated Scale of Intelligence (Wechsler, 2014) was 109.7 (SD = 10.9, range = 82-141). IQ scores were not significantly correlated with age (r =. 19, p =. 076), nor did IQ significantly differ between males (mean = 111.1, SD = 12.1) and females (mean = 108.4, SD = 9.5, t = 1.2, p =. 236).

### Stimuli and task

EEG was recorded during a three-stimulus visual oddball paradigm (**Fig.1**). The task was administered using the E-Prime software and presented on a 19-inch computer screen with a viewing distance of approximately 80 cm, and responses were obtained on a PST Serial Response Box. Stimuli were presented on a black screen for 1 s, with a 2 s interstimulus interval during which a white fixation cross was presented. Target and standard stimuli were both blue ellipses, with a height and width of 17.5 x 14.5 cm for the target and 15 x 12 cm for the standard stimulus. The distractor stimulus was a large blue rectangle of 21 x 17 cm. The resulting visual field was about 9° × 7°, 10° × 8°, and 12° × 10° for the standard, target, and distractor stimuli, respectively. Participants were told that they would be shown small and large blue circles and instructed to respond by key press when presented with the large circle, and not respond to any other stimuli. All participants first completed a short practice block consisting of 10 trials (7 standard, 3 target). The instructions were then repeated before participants completed the main task block, which consisted of a total of 250 stimuli presented in pseudorandomized order (200 standard, 25 target, 25 distractor). Data sets were included in the final analysis when a minimum of 75% hits on target trials and a maximum of 30% responses to standards or distractors were scored. Two participants were excluded due to having too few hits. This experimental paradigm is a variation of one used by Comerchero and Polich (1999), which has been shown to elicit both P3b and P3a event-related potentials (ERPs) (Polich, 2004), and which we have used in previous developmental and aging studies (Fjell, Rosquist, & Walhovd, 2009; Fjell et al., 2007; Stige et al., 2007).

**Fig. 1.**
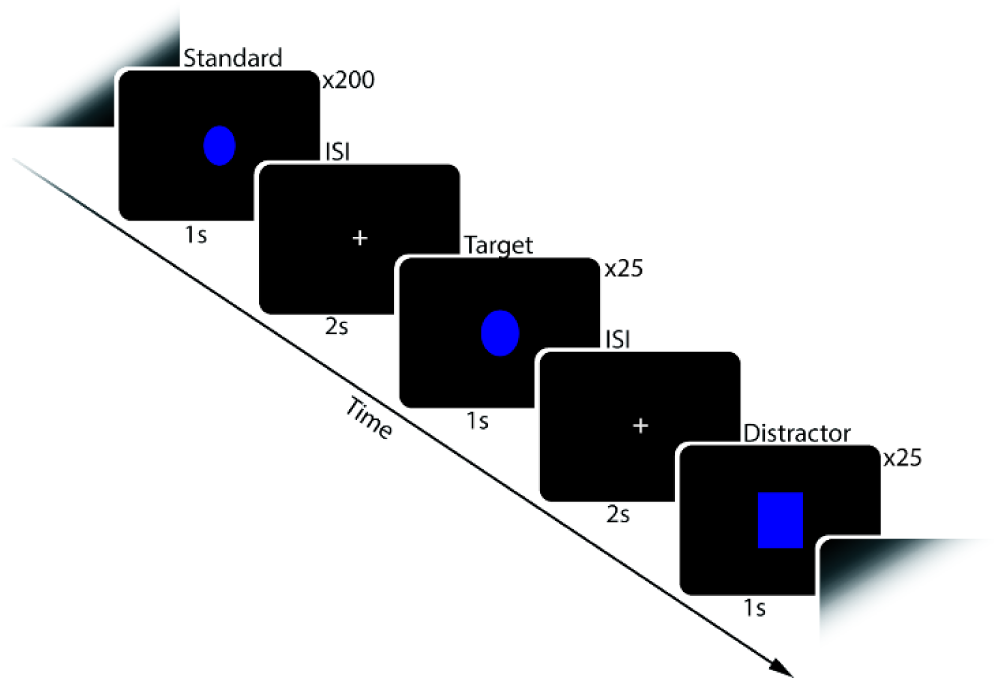
Schematic illustration of the visual oddball task. Participants were asked to respond by button presses to target stimuli (25 ellipses) and ignore other stimuli. These included standard stimuli (200 smaller ellipses), as well as distractor stimuli (25 rectangles).

### EEG data acquisition

Participants performed the task in an electrically shielded room while seated in a comfortable high-back chair. The electrophysiological recordings were done using 128 EEG channels with an electrode placement according to the 10% system (EasyCap Montage No. 15, http://www.easycap.de/). The sampling rate during recording was set to 1000 Hz. The electrodes used were EasyCap active ring electrodes (Ag/AgCl) with impedance conversion circuits integrated into the electrode housing that allows high quality recordings even with high electrode impedance values, thus reducing preparation time and noise. The signals were amplified via a Neuroscan SynAmps2 system and filtered online with a 40 Hz low-pass and a 0.15 Hz high-pass analog filter prior to digitization and saving of the continuous data set. During recording, all electrodes were referenced to an electrode placed on the left mastoid. Vertical eye blinks were recorded with one electrode above and one electrode below the left eye, and a ground electrode was placed anteriorly on the midline.

### EEG data processing

Initial pre-processing of the EEG data was done using the Curry 7 neuroimaging suite (http://compumedicsneuroscan.com/curry-neuroimaging-suite/). The continuous EEG data of each subject was visually inspected to identify noisy electrodes, which were then interpolated as the mean of its 8 nearest neighboring electrodes (excluding noisy neighboring electrodes). Data from 12 participants was excluded due to noise that could not be corrected through this interpolation procedure. This included data where noisy electrodes were clustered together, making interpolation from clean electrodes impossible, as well as cases where large portions of the data contained movement artifacts. The continuous EEG was then filtered from 1Hz to 20Hz. Eye-blink artifacts were corrected for using a regression-based approach as implemented in CURRY. The data sets were then segmented into 900 ms epochs, starting 100 ms before each stimulus presentation. Epochs were baseline corrected relative to the 100 ms time window preceding stimulus presentation.

Group-level blind source separation was applied to estimate a single set of components that capture the representative activity from neural sources commonly expressed across the whole sample (Huster et al. 2015). To do so, an equal number of trials was extracted from each participant. 19 trials from the target condition, 19 trials from the distractor condition, and 38 from the standard condition were randomly selected for each subject. Only trials with correct responses to targets and without false positive responses to distractor or standard stimuli were included in the analysis. The group decomposition procedure requires the same number of trials per condition and subject, and thus the number of trials extracted per condition was determined based on the number of correct trials that further survived artifact correction. In addition, the number of standard trials was chosen to match the total number of distractor and target trials to avoid biases in the estimation procedure. Then, group-SOBI (second order blind identification; Belouchrani, Abed-Meraim, Cardoso, & Moulines, 1997) was applied to the EEG, preceded by two consecutive principal component analysis steps (PCA; single subject and group-level) as a means of data reduction (Eichele et al. 2011; Huster et al. 2015). Firstly, for each participant, all trials were vertically concatenated and z-standardized over channels, resulting in a two-dimensional matrix with electrodes as rows and time-points as columns spanning all 76 selected trials. This matrix was then entered into a subject-specific PCA. 4 PCs were selected from each participant, on average accounting for 70% of the variance in the single-subject data. 70% was chosen as a threshold because higher cutoffs resulted only in more noisy components without changing the main, stable components. These 4 components were extracted from each subject’s data, concatenated vertically (along the dimension originally representing channels), and entered into a second, group-level PCA. Again, 4 group-PCs were extracted and subjected to SOBI (with temporal lags up to 50 data points or 100 ms), resulting in 4 group-level component time-courses. Finally, in order to examine between-subject differences in the group-level components, single-subject activation of those components were reconstructed by matrix multiplication of the original single-subject EEG time-courses with the demixing matrices resulting from the two preceding PCA’s and SOBI. (see Eichele et al. 2011; Huster et al. 2015; Huster & Raud, 2018 for details).

We then selected components for further analysis best capturing those ERPs clearly elicited in this paradigm: N1, P2, P3a and P3b. These were identified by visually inspecting the component scalp distributions and the timing of component peaks. This led to the selection of three components (see below). In each of these components, we identified peaks of interest and then extracted the area under the curve from a 40 ms time window surrounding each peak of interest for each experimental condition for further statistical analysis (hereafter referred to as time frames). The summed area under the curve in each time frame will hereafter be referred to as that component’s strength. This measure of component strength is conceptually somewhat similar to ERP amplitude. However, a component’s strength is in arbitrary units and the direction of the component deflections does not necessarily reflect the charge of the amplitudes in the EEG data.

### MRI Acquisition

MRI data were acquired using a 1.5 Tesla Siemens Avanto scanner (Siemens Medical Solutions) with a 12-channel head coil. For the morphometric analyses we used a 3D T1-weighted MPRAGE pulse sequence with the following parameters: TR/TE/TI/FA = 2400 ms/3.61 ms/1000 ms/8°, matrix 192 × 192, field of view = 240, 160 sagittal slices, voxel size 1.25 × 1.25 × 1.20 mm. Duration of the sequence was 7 min 42 s. A minimum of two repeated T1- weighthed sequences were acquired. All images were screened immediately after data acquisition and rescanning was performed if needed and possible. The protocol also included a 176-slice sagittal 3D T2-weighted turbo spin-echo sequence (TR/TE = 3390/388 ms) and a 25-slice coronal FLAIR sequence (TR/TE = 7000–9000/109 ms) to aid the radiological examination.

### MRI Data Processing

For each participant, the T1-weighthed sequence with best quality as determined by visual inspection of the raw data was chosen for further analysis. Whole-brain volumetric segmentation and cortical reconstruction was performed with FreeSurfer 5.3, an open source software suite (http://surfer.nmr.mgh.harvard.edu/). The details of the procedures are described elsewhere (Dale, Fischl, & Sereno, 1999; Fischl, 2012; Fischl et al., 2002; Fischl, Sereno, & Dale, 1999). Briefly, the processing includes motion correction, removal of non-brain tissue, automated Talairach transformation, segmentation of structures, intensity normalization, tessellation of surfaces, automated topology correction, and surface deformation to optimally place tissue borders. Cortical surface area (white matter surface) maps were computed by calculating the area of every triangle in the tessellation. The triangular area at each location in native space was compared with the area of the analogous location in registered space to give an estimate of expansion or contraction continuously along the surface (“local arealization”) (Fischl et al., 1999). Cortical thickness maps for each subject were obtained by calculating the distance between the cortical gray matter and white matter surface at each vertex (Fischl & Dale, 2000). The maps produced are not restricted to the voxel resolution of the original data and are thus capable of detecting submillimeter differences. All processed scans were visually inspected in detail for movement and other artifacts. Minor manual edits were performed by trained operators on 8 subjects, usually restricted to removal of non-brain tissue included within the cortical boundary. All scans were deemed sufficiently free of movement noise to be included. Before statistical analyses, the surface maps for cortical area and thickness were smoothed with a Gaussian kernel of full-width at half maximum of 15 mm.

### Statistical Analysis

Descriptive statistics and Pearson’s correlation were used to characterize the sample and task performance, and to test how task performance was associated with age. Pearson’s correlation coefficients were computed to assess the relationships between a component’s strength and age. Partial correlations were computed between a component’s strength and task performance, controlling for age, to rule out task improvement due to maturation in general and thus examine individual differences in the maturational processes indexes by the measured components. Surface-based cortical analyses were performed vertex-wise (point-by-point) using general linear models, as implemented in FreeSurfer 5.3. Main effects of the strength of each extracted component on cortical structure was tested, while controlling for the effects of sex and age. Separate analyses were performed for cortical surface area and thickness maps. The data were tested against an empirical null distribution of maximum cluster size across 10,000 iterations using Z Monte Carlo simulations as implemented in FreeSurfer (Hagler, Saygin, & Sereno, 2006; Hayasaka & Nichols, 2003) synthesized with a cluster-forming threshold of P < .001, yielding clusters corrected for multiple comparisons across the surfaces. Cluster-wise corrected P < .01 was regarded significant. Mean vertex-wise cortical surface area or thickness values were then extracted from each significant cluster.

## RESULTS

### Task performance

Participants responded correctly to an average of 94% (SD =. 07, range =. 76-1.00) of targets with an average median reaction time of 669 ms (SD = 111, range = 346-989). Standard and distractor stimuli were incorrectly responded to 2% (SD =. 04, range =. 00-.29) and 1% (SD =. 02, range =. 00-.12) of the time, respectively, and with average median reaction times of 780 ms (SD = 192 range = 346-1377) and 591 ms (SD = 127, range = 439-881), respectively. The number of correct responses (hits) was positively correlated with age (r =. 28, p =. 009), while the number of false positives by responding to standard stimuli, but not to distractor stimuli, was negatively correlated with age (r = -.30, p =. 004 and r =. 047 p=. 669). Reaction times for correct responses to target and incorrect responses to both standard and distractors were significantly related to age, with older participants having faster reaction times (Target: r = -.62, p < .001; Standard: r = -.33, p =. 009; Distractor: r = -.49, p =. 010).

### EEG decomposition

The SOBI decomposition of the EEG epochs yielded four components (**Fig.2**). We visually selected those three components that best reflected early visual or P300-like processing as inferred from both component topographies and activity patterns. A total of four peaks of interest were identified for components 2 through 4, with peak latency defined separately for each experimental condition. These were the early positive and the early negative peaks of component 2 (target: 132 ms/216 ms, distractor: 132 ms/222 ms, standard: 133 ms/216 ms), likely representing the visual N1 and P2; the second positive peak of component 3 (target: 346 ms, distractor: 304 ms, standard: 312 ms), presumably representing a fronto-central P300; the large positive peak of component 4 (target: 464 ms, distractor: 428 ms, standard: 328 ms), likely represents a parietal P300. The assumptions about the nature of each component are based on their locations on the scalp, the degree of difference between experimental conditions, and the similarity in timing and appearance to the ERPs expected from the experimental paradigm. The sign of each SOBI component peak is not indicative of the amplitude charge in the raw data before the decomposition. The actual amplitude of a component after its back-projection to the EEG corresponds to the product of the component weight at a given electrode and its activity. For instance, if an electrode’s weight is negative the EEG potential will be positive if the component activity is negative as well, whereas it will be negative if the component activity is positive. For each peak of interest, the area under the curve +/-20 ms from the peak was extracted, separately for each condition. From this point forward the terms fronto-central P300, parietal P300, N1 and P2 will be used to describe the extracted component time frames. The term strength will be used for the summed values of the extracted time frames for each component.

**Fig. 2.**
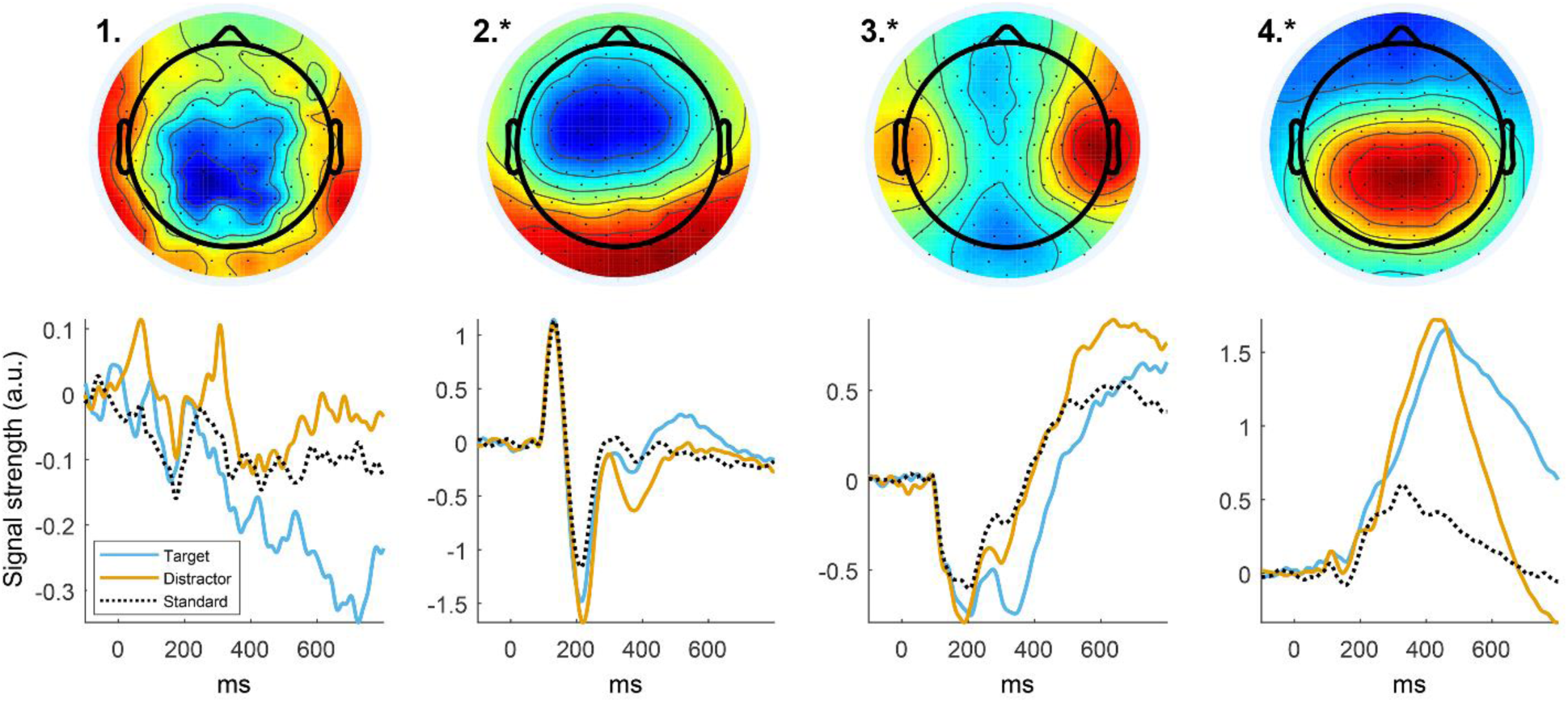
Component output of Group level SOBI decomposition of all EEG epochs. Top: Component scalp topographies. Bottom: ERPs of all independent components for standard (black, dotted), target (blue) and distractor (orange) conditions. Components marked with (*) were kept for further analyses. Peaks of the rightmost three components are assumed to represent the following: (2) N1 and P2; (3) fronto-central P300; (4) Parietal P300. From these we extracted the area under the curve in a 40 ms time window, which were used for all further statistical analyses.

### Age-related differences in EEG component strength

Correlation analyses were performed to investigate the associations between the strength of each of the EEG components and age (**Fig.3**). There were significant positive correlations between age and the parietal P300 for the target condition (r =. 24, p =. 029) and the distractor condition (r =. 34, p =. 001), with component strength increasing concurrently with age. No significant association was found for the standard condition (r = -.16, p =. 149). Significant negative correlations were found between age and the N1 for all conditions (Target: r = -.42, p < .001; Distractor: r = -.39, p < .001; Standard: r = -.35, p =. 001), indicating a decrease in strength with age. Significant positive correlations were found between age and the strength of the P2 for all experimental conditions, corresponding to decreasing component strength with age (Target: r =. 40, p < .001; Distractor: r =. 28, p =. 009; Standard: r =. 45, p < .001). There were no significant associations between age and the fronto-central P300 in any of the conditions (Target: r = -.02, p =. 866; Distractor: r = -.05, p =. 662; Standard r = -.08, p =. 483). In sum, the results indicate that the parietal P300 components get stronger with age during development, while the N1 and P2 get weaker.

**Fig. 3.**
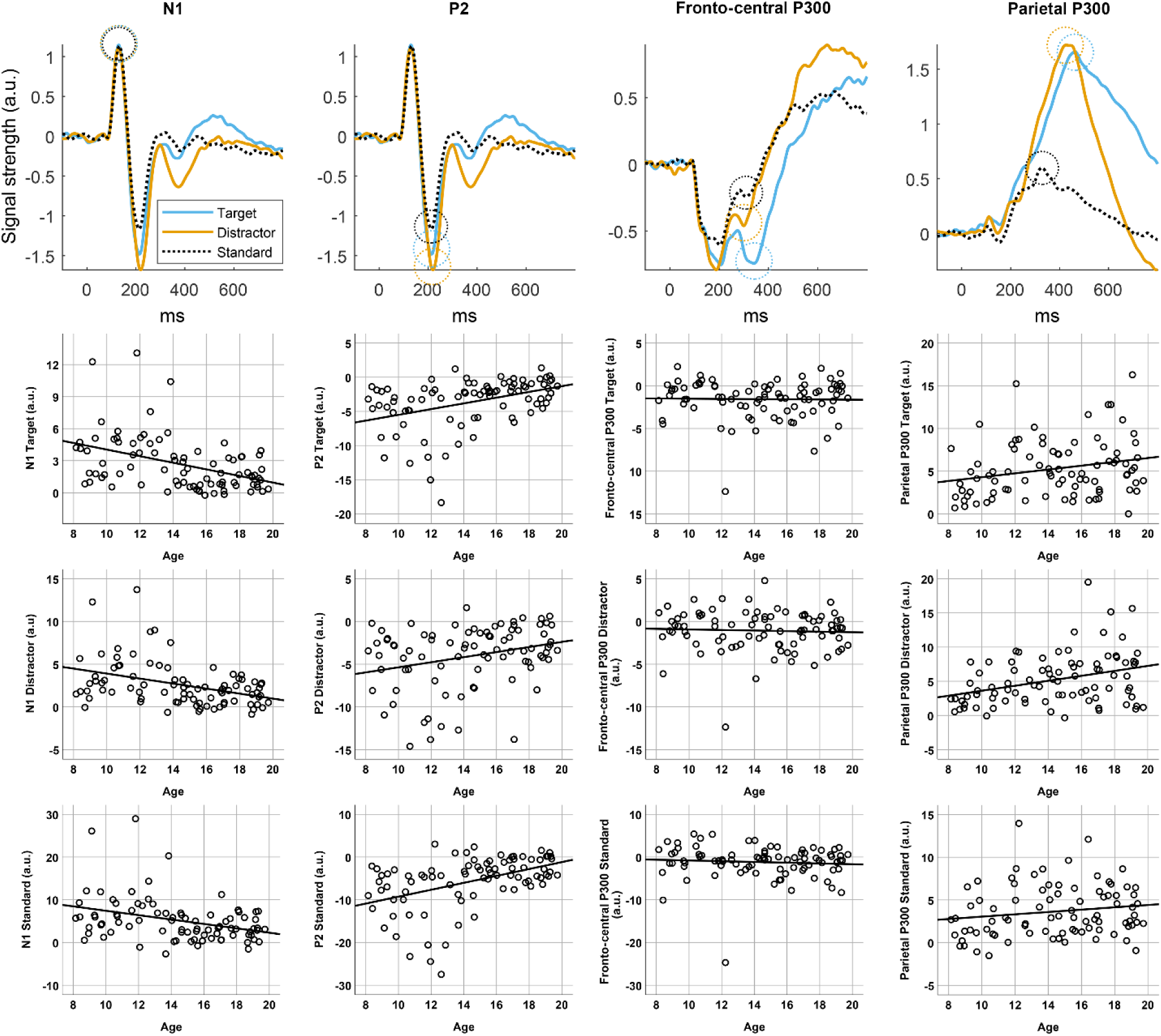
Age-related differences in EEG component strength. Top: ERPs of the three SOBI components used in our analyses for standard (black, dotted), target (blue) and distractor (orange) conditions. 40 ms time windows of each extracted peak area for each experimental condition are marked with dotted circles. Signal strength is plotted in arbitrary units, and the signs of these values do not necessarily reflect the charge of the EEG amplitudes in the data before the decomposition. Bottom: Scatter plots showing relationships between age and component strength for the extracted time windows of each component.

### Associations between EEG component strength and task performance

Partial correlations were performed to test the associations between the strength of all EEG components and task performance variables, including hits on targets, false positives for standard and distractor stimuli, and median reaction times, with age included as a covariate. The strength of the standard and distractor conditions for the parietal P300 component were positively associated with number of correct responses (hits) (Distractor: r =. 21, p =. 050; Standard; r =. 25, p =. 024), but this was not the case for the target condition (r =. 11, p =. 310). The strength of the fronto-central P300 showed a significant negative association with the number of hits for the target and distractor conditions (Target: r = -.25, p = -.021; Distractor: r = -.23, p =. 032), indicating better precision with a stronger fronto-central P300 component, but not for the standard conditions (r = -.14, p =. 217). Median reaction time to targets was positively associated with the strength of the fronto-central P300 for the standard condition (r =.27, p =. 013), as well as the strength of P2 for standard (r =. 24, p =. 030) and distractor (r =. 28, p =. 010) conditions.

### Associations between EEG component strength and cortical structure

Finally, to examine the associations between the strength of the EEG components and cortical structure, surface-based cortical analyses were performed using GLMs, controlling for sex and age. Separate analyses were performed for cortical surface area and cortical thickness. After correction for multiple comparisons using cluster size inference, the results revealed a cluster in the inferior temporal gyrus of the left hemisphere where a stronger parietal P300 in both the target condition (cluster size = 519.28 mm^2^, clusterwise p =. 006, MNI max vertex (X, Y, Z) = -49.4, -18.2, -34.7) and the distractor condition (cluster size = 705.38 mm^2^, clusterwise p =. 001, MNI max vertex (X, Y, Z) = -45.5, -16.0, -34.8) were significantly associated with larger cortical surface area (**Fig.4**). No other effects reached significance after controlling for multiple comparisons.

**Fig. 4.**
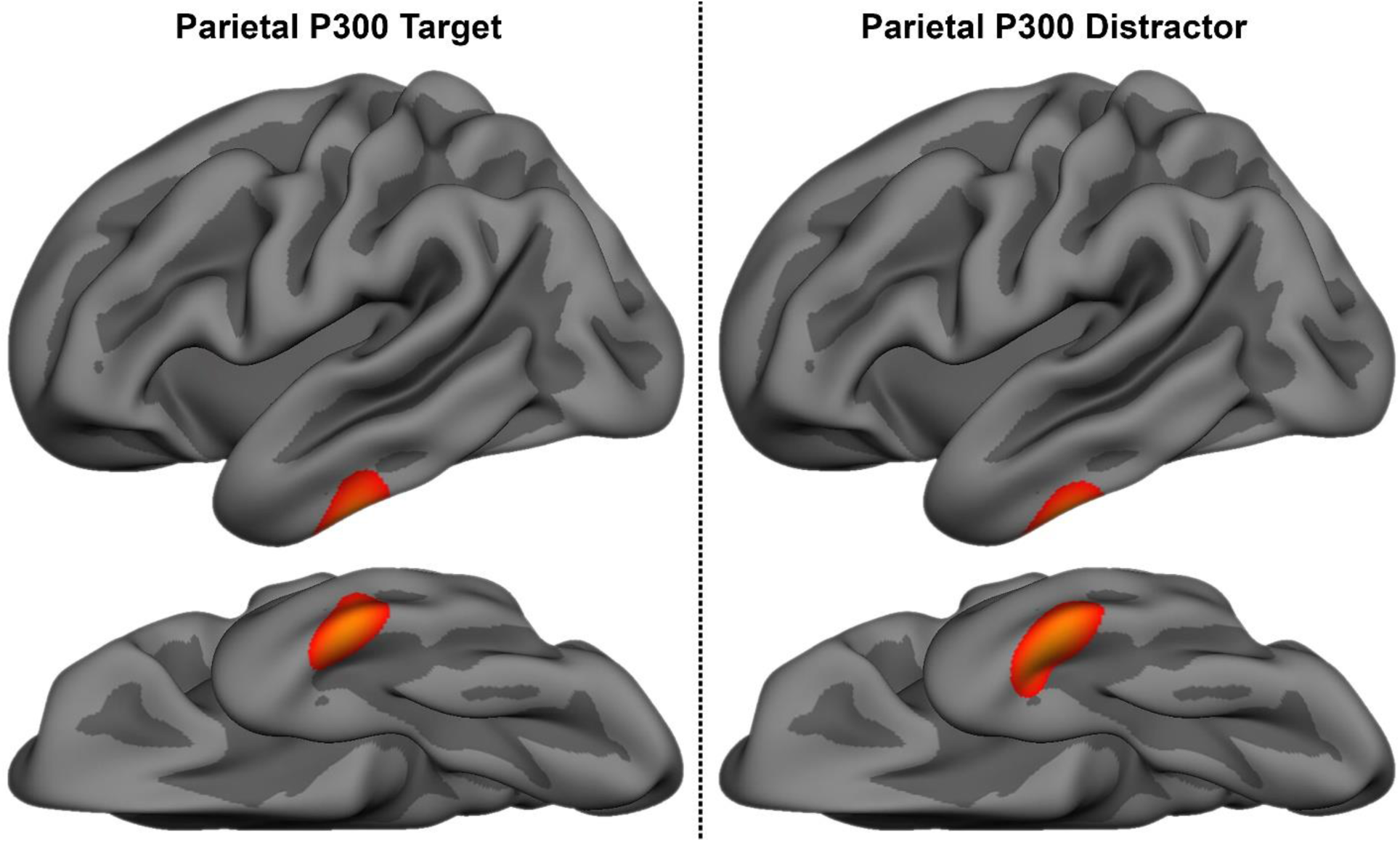
Associations between EEG component strength and cortical structure. Clusters in the left hemisphere at cluster-forming thresholds of P < .001 showing a significant association between cortical surface area and parietal P300 activation, independently of age and sex for the target (left) and distractor (right) conditions.

## DISCUSSION

In this study, we investigated age-related differences in early visual and P300 components, and how these components are related to task performance and cortical surface area and thickness. Group-level blind source separation identified three components of interest. Based on the component topographies and temporal activity profiles, they were deemed to represent a parietal P300, a fronto-central P300, as well as the visual N1 and P2. N1 and P2 decreased in strength with age; while in contrast, the strength of the parietal P300 increased with age. Individual differences in the strength of the P300 components were positively associated with aspects of better task performance, while the P2 showed associations in the opposite direction. The strength of the parietal P300 component was positively associated with cortical surface area in the left lateral inferior temporal lobe. The associations between the EEG components and age, task performance and cortical structure are discussed further below.

### Relationships of age to the identified electrophysiological components

We identified two P300-like components: a fronto-central P300 and a parietal P300. The parietal P300 component increased in strength with increasing age. This result is in accordance with those of van Dinteren and colleagues’ large cross-sectional study and meta-analysis (van Dinteren et al., 2014a), where auditory P300 amplitudes were found to increase through childhood and adolescence and peak in the early 20s. However, the literature on the development of the visual P300 is relatively sparse, and the developmental trajectory is less clear, with several discrepant findings. It has been hypothesized that the developmental trajectory of the P300 might differ for auditory and visual modalities (Segalowitz et al., 2010), since some studies using visual oddball paradigms have found negative age associations. A study of adolescent girls aged 14-19 found the visual P3b to decrease with age in a control group, but not in a clinical group with borderline personality disorder (Houston, Ceballos, Hesselbrock, & Bauer, 2005). Similarly, another study indicated that the visual P3b decreases with age in participants 8-18 years (Hill & Shen, 2002). Stige et al. (2007) also found visual P3a and P3b to decrease from late childhood to adolescence. All studies mentioned used variants of visual oddball tasks, and it is not clear why these studies found opposite results to the present study. However, developmental studies on visual P300 are still rare and several of the available studies included small samples. To test whether P300 development differs between modalities, future studies will have to directly compare age-related differences in P300 components using similar paradigms with the same participants. Unlike the parietal P300, the fronto-central P300 component did not show any age-related changes. The reason for this difference is unclear; as is the functional distinction between the two observed P300 components. A common interpretation of the P300 is that it reflects multiple overlapping signals from several sources (Bocquillon et al., 2011; Wronka et al., 2012), with slightly different functions for the fronto-central and the more parietal P300. The earlier peak and more anterior topography of the fronto-central P300 could indicate a similarity with the P3a, while the parietal P300 component seems more similar to the somewhat later and more posterior P3b. If this speculative interpretation is correct, the lack of a correlation with age would suggest that the P3a, which is tied to stimulus-driven frontal attention mechanisms during task processing (Polich, 2007), has already reached maturity in late childhood. This is in line with the limited research on the development of P3a (Segalowitz et al., 2010). Following this reasoning our results would indicate that the P3b, which is associated with attention and subsequent memory processing (Polich, 2007), shows a more protracted development.

Our decomposition of the EEG data also identified an early visual component which had two marked peaks. These were similar to, and likely reflect, the N1 and P2 evoked potentials. The N1 is thought to reflect activity tied to controlled information processing (Segalowitz et al., 2010), while the P2 is tied to working memory function (Finnigan, O’Connell, Cummins, Broughton, & Robertson, 2011). In contrast to the parietal P300 component, which increased in strength with age, our results showed that both of these early visual components decreased in strength with increasing age. This is consistent with previous developmental studies on N1 (Itier & Taylor, 2004; Tomé, Barbosa, Nowak, & Marques-Teixeira, 2015) and P2 (Ponton, Eggermont, Kwong, & Don, 2000). A possible confounding factor is developmental increase in skull density and thickness, which could also explain a reduction in EEG signal strength through development. However, this likely only has a minor impact across adolescence (Segalowitz et al., 2010), and so is unlikely to explain the age correlations observed in our sample, particularly considering the positive age associations observed for the parietal P300 component, though it might have a modulating influence.

### Relationships of the identified electrophysiological components and task performance

The strength of N1 was not associated with task performance in our data. This does not lend support to the notion proposed by Segalowitz et al. (2010) that lowering of the N1 amplitude might reflect a narrowing of attentional focus and increased vigilance. However, lower P2 strength was associated with faster reaction times in distractor and standard conditions, and N1 and P2 are commonly viewed as highly dependent parts of the visual evoked potential, though they are not functionally identical (Crowley & Colrain, 2004).The dissociation seen in the age-related differences for the parietal P300 component in contrast to the early visual components was also reflected in the direction of the associations we found between the strength of the EEG components and aspects of task performance. In contrast to the negative association in P2, the strength of the parietal P300 component was positively associated with aspects of task performance. The strength of the parietal P300 in the standard and distractor conditions was associated with number of correct responses. Also, a stronger fronto-central P300 in the target and distractor conditions was associated with a better hit rate and in the standard condition with faster reaction times. Positive associations between P300 amplitudes and oddball task performance were reported by van Dinteren et al. (2014b), and several developmental studies using stop signal tasks have shown successful trials to be associated with higher P300 amplitudes compared to unsuccessful ones (Dimoska, Johnstone, & Barry, 2006; Dimoska, Johnstone, Barry, & Clarke, 2003; Overtoom et al., 2002). As we controlled for age in our analyses, these results suggest that better task performance was related to EEG signatures more similar to older, more mature adolescents. This might indicate that the components reflect individual differences in maturation of the visual attentional system beyond what can be explained by age alone. In sum, N1 and P2 strength was lower for older adolescents and lower P2 strength was associated with better task performance, while the opposite pattern was generally seen for the P30 components. From this we speculate that improvements in attention seen in adolescence stems partly from the attentional system becoming more efficient at distributing cognitive resources to task relevant attentional processes. The relative shift from early visual to P300 activity could correspond to a shift in task strategy or the development of cognitive resources. As the P300 is thought to be associated with complex attentional mechanisms such as stimulus comparisons and the facilitation of memory processing (Polich, 2007), this could reflect a shift away from the mere reliance on early and quick feature detection and towards a more efficient comparative categorization approach.

### Relationships between the identified electrophysiological components and cortical characteristics

Using gaussian-based Monte Carlo simulations in surface-based clusterwise analysis has been shown to underestimate the false positive rate (Greve & Fischl, 2017). To correct for this, we used a high smoothing level (FWHM 15 mm), and a strict cluster-forming threshold of P < .001. We found an age-and sex-independent positive association between the strength of the parietal P300 and cortical surface area in the left inferior temporal gyrus for both the target and distractor conditions. This is generally consistent with other studies of both children and adults, indicating positive associations between various cognitive functions and cortical surface area (see e.g. Curley et al., 2017; Fjell et al., 2012; Walhovd et al., 2016). No associations were found between the strength of the fronto-central P300 or the early visual component and cortical surface area, or between any of the EEG components and cortical thickness. Controlling for age was done to prevent potenially specific associations between P300 strength and cortical structure from being masked by the widespread age-related structural changes in the age-range studied (Tamnes et al., 2017). After controlling for age, the findings can be interpreted as the relative association between cortical structure and P300 strength within each age group.

Multiple brain regions are known to contribute to P300 generation (Friedman, 2003). Intriguingly, the inferior temporal gyrus has previously been implicated in generating the visual P3b. A combined fMRI/EEG study by Bledowski et al. (2004) that used a three-stimulus visual oddball task similar to the one used in our study found P3b to be generated by inferior temporal and parietal regions, while P3a was found to be produced mainly by the insula and frontal regions. The authors suggest that the cortical activation in the inferior temporal region during P3b generation could reflect the categorization of visual stimuli by higher visual areas. However, other areas of the temporal lobes have been pegged as possible sources of the P300 more generally, along with several other regions. Several neural generators of the auditory P300 have been suggested, including the prefrontal cortex, the temporo-parietal junction and the primary auditory cortex (Friedman, 2003). A meta-analysis of fMRI studies using visual and auditory oddball tasks by Kim (2014) concluded that oddball effects were strongly associated with a ventral, modality-independent attentional network. This network included the temporo-parietal junction, the anterior medial frontal gyrus, the anterior insula and the anterior cingulate cortex, as well as the inferior frontal junction and the sensory cortical regions associated with the sensory modality relevant for the task. In a source localization study of the P300 using a visual oddball paradigm similar to the one used in our study, Bocquillon et al. (2011) suggested that the dorsal frontoparietal network generates the P300, with inferior parietal areas involved in target-specific processing. Of studies on cortical structure, Fjell et al. (2007) found that higher P3a amplitude was positively related to cortical thickness in parietal regions, the temporo-parietal junction and in the middle frontal gyrus, but only for elderly adults.

In sum, the inferior temporal gyrus region that was found to be associated with the strength of the parietal P300 does not overlap with previously identified modality independent P300 sources but is known to be involved in visual shape processing (Gerlach et al., 2002) and has been identified as one source of the visual P3b by Bledowski et al. (2004). From this we can speculate that the observed association between larger cortical surface area in this region and greater parietal P300 strength is indicative of a maturational difference in attentional target processing in the visual modality. Longitudinal studies are however needed to clearly establish developmental trajectories of P300 components and how these changes relate to structural brain maturation. In addition, methodological studies comparing the results of studies using decomposition techniques and traditional ERP analyses for well-studied samples and paradigms could prove enlightening.

## Conclusions

Our results suggest that the parietal P300 increases in strength with increasing age through adolescence, while early visual component strength decreases. Task performance improved with age and was associated with stronger P300 components and a weaker N2 component, independently of age. Finally, parietal P300 strength was, independently of age, associated with larger cortical surface area in inferior temporal gyrus for the left hemisphere. We suggest that the results reflect functional brain maturation in terms of increasing brain responses to task-relevant stimuli, reflective of a more mature and efficient visual attention system.

Conflict of Interest: The authors declare that they have no conflict of interest.

Ethical approval: “All procedures performed in studies involving human participants were in accordance with the ethical standards of the institutional and/or national research committee and with the 1964 Helsinki declaration and its later amendments or comparable ethical standards.”

## Acknowledgments

This study was supported by the Research Council of Norway (to KBW and 230345 to CKT), and the University of Oslo (to KO).

